# Exploring Genomic Conservation in Actinobacteriophages with Small Genomes

**DOI:** 10.1101/2022.02.05.479200

**Authors:** Vincent Y. Tse, Haoyuan Liu, Andrew Kapinos, Canela Torres, Breanna Camille S. Cayabyab, Sarah N. Fett, Lucy G. Nakashima, Mujtahid Rahman, Aida S. Vargas, Krisanavane Reddi, Jordan Moberg Parker, Amanda C. Freise

**Author notes:** Corresponding Author: Amanda C. Freise.

## Abstract

Actinobacteriophages of a wide range of genome sizes continue to be isolated and characterized, but only a handful of these have atypically small genomes, defined in this work as genome sizes under 20,000 bp. These “small phages” are relatively rare and have received minimal study thus far. Of the actinobacteriophages published in PhagesDB.org, small phages have been isolated on *Arthrobacter*, *Gordonia*, *Rhodococcus*, and *Microbacterium* hosts. A previous study by Pope *et al*. showed that *Gordonia* small phages have similar gene products and amino acid sequences. Here, we set out to further examine relationships between small *Gordonia* phages as well as small phages that infect other hosts. Of the 3222 sequenced phages listed on PhagesDB, we identified 109 distinctly small phages with genome sizes under 20,000 bp. The majority of the small phages were isolated on *Arthrobacter* or *Microbacterium* hosts. Using comparative genomics, we searched for patterns of similarity among 34 cluster-representative small phages. Dot plot comparisons showed that there was more amino acid conservation than nucleotide identity amongst small phages. Gene content similarity (GCS) analysis revealed that the temperate *Gordonia* phages in Cluster CW share significant GCS values (over 35%) with the lytic *Arthrobacter* phages in Cluster AN, suggesting that some small phages have a considerable degree of genomic similarity with each other. SplitsTree analyses of shared phams (genes with substantial amino acid identity) supported the complexity of clustering criteria in small phages, given shuffling of genes across phages of different clusters and close relationships despite varied cluster membership. We observed this continuum of phage diversity through *Rhodococcus* phage RRH1’s closer similarity to phages in *Gordonia* subcluster CW1 than CW1 is to *Gordonia* subcluster CW3. Finally, we were able to confirm the presence of conserved phams across not only small *Gordonia* phages but also within small phages from different clusters and hosts. Studying these genomic trends hidden in small phages allows us to better understand and appreciate the overall diversity of phages.

## Introduction

Bacteriophages have been described as among the planet’s most influential and are the most abundant biological entities on Earth, numbering up to an estimated 10^31^ individual phage particles and playing key roles in the environment [1,2]. The enormous diversity of phages has led to observations of phages that infect a wide variety of hosts, exhibit an array of life cycles, and display an assortment of genomes [3]. Over the years, phages of many different varieties continue to be isolated and added to PhagesDB, a database for actinobacteriophage research [4]. At the time of our study, we observed a sizable gap in the distribution of genome sizes on PhagesDB, wherein actinobacteriophages with genome sizes nearing 20,000 base pairs were separated from the rest of the phage database. As such, we ultimately designated these phages under 20,000 base pairs as atypically small phages. We have identified 109 small actinobacteriophages and characterized 34 of those phages as representative small phages. Few phages of this size have been analyzed, but due to their simplicity, understanding the overarching structure of these genomes can provide a solid foundation for contextualizing future phage research [5].

The goal of this study was to characterize these phages with smaller genomes and shed a clear light on patterns of genomic conservation that have previously been obscure. Bacteriophage genomes have often been shown to display a mosaic nature [6] and a continuum of diversity [7]; as atypically small phages have a very compact and simple genomic structure, we wanted to learn whether small phages share some of these very few genes with each other. A recent study showed that atypically small *Gordonia* phages have similar gene products and amino acid sequences [8] but whether these *Gordonia* small phages display similarity with other small phages from different hosts and clusters was not clear. Another subsequent study likewise suggested that very small *Microbacterium* phages share similar genome architecture with small *Arthrobacter*, *Gordonia*, and *Rhodococcus* phages [9]. Further analysis of these phages, particularly of newly isolated small phages, can help reinforce genomic patterns that have already been observed and even identify new trends that may have been difficult to see.

In this study, we have identified and characterized atypically small actinobacteriophages. Through dot plot analysis, gene content similarity comparisons, and the construction of a Splitstree network phylogeny, we examined relationships between these small phages and were able to support the findings presented in the aforementioned studies. We additionally observed the conservation of several phams among otherwise diverse groups of small phages, such as genomic similarity between *Arthrobacter* phages, *Gordonia* phages, and RRH1, a *Rhodococcus* phage that is currently the smallest phage in the Actinobacteriophage database [5]. Overall, this indicates that genome mosaicism is prevalent even in small phages with few genes and supports the presence of a continuum of phage diversity.

## Methods

### Phage preparation, DNA extraction, and genome sequencing and annotation

Various small genome phages were isolated from soil samples and phage DNA was both purified and amplified according to the procedures described by SEA-PHAGES [10,11]. The phage genomes were then sequenced using Illumina-MiSeq and assembled as previously done [12] and auto-annotated using DNA Master [13], Glimmer [14], and GeneMark [15]. Various programs were used to manually verify the accuracy of the auto-annotation, such as Starterator [16], PhagesDB and NCBI BLASTp [4,17], NCBI Conserved Domain Database [18,19], and HHpred [20].

### Comparative genomic analyses

A list of all 3222 phages and their genomic data as of November 2020 was obtained from PhagesDB (https://phagesdb.org/data/). The phages were plotted by increasing genome size. In order to determine the genome size cutoff for “small” phages, we identified a large gap in genome size from 19,679 bp to 28,876 bp and thus set 19,679 bp as the cutoff for “small” phages. Small phages were subsequently sorted based on their isolation type, host, morphotype, life cycle, GC content, and cluster. Representative small phages (Table 1) were selected such that as much diversity as possible was maintained from the aforementioned categories.

**Table 1.**
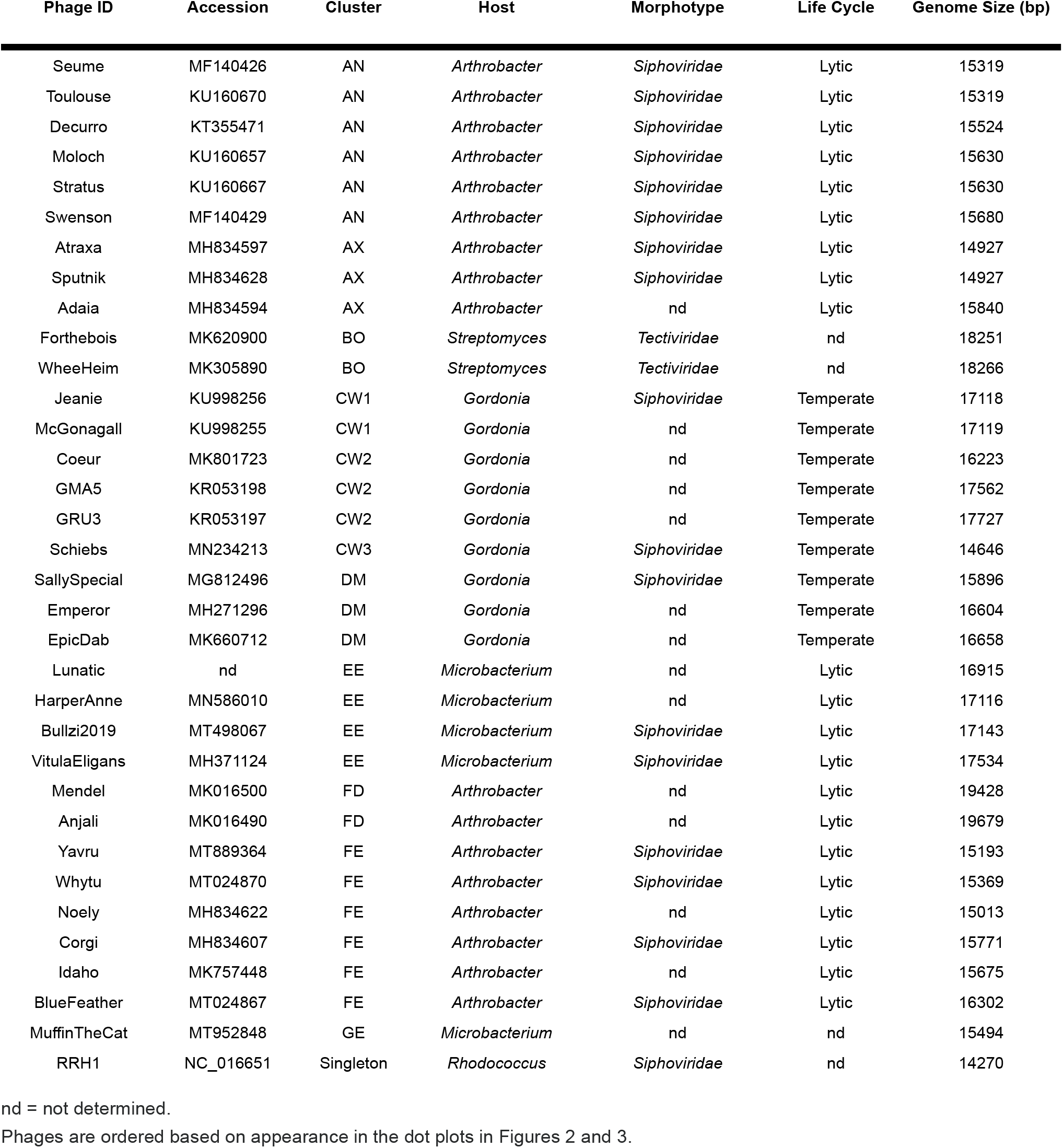
34 representative phages from each cluster of small genome phages show immense physical and genomic diversity.

Nucleotide and amino acid FASTA files from each representative small phage were extracted from NCBI and imported into Gepard 1.40 [21] to produce dot plots. Furthermore, gene content similarity (GCS) values of the 34 representative phages were computed using the PhagesDB Explore Gene Content tool [4]. The GCS values of all 34 phages were entered into GraphPad Prism9 to generate a heatmap. Pham data for all representative small phages was extracted from the Actino_Draft database (version 382) and imported into SplitsTree 4.16.2 [22]. Using default parameters, a network phylogeny was created for these representative small phages.

To expedite the process of genome structure comparison, 11 of the 34 representative phages were chosen in the same fashion as how the initial 34 were selected from the original 109 phages. These 11 representative phages were used to construct a comparative pham map. Genome maps for each of the 11 phages were downloaded on Phamerator and annotated on Inkscape 1.0 (https://inkscape.org/).

## Results

### 109 bacteriophages were designated as “small genome phages” and 34 representative phages were selected for analysis

To determine the genome size cut-off as the upper limit of “small” phages, a size continuum graph was constructed to list all the phages from PhagesDB in order of increasing genome size (Figure 1A). A large gap in genome size was observed between the 109th and 110th smallest phage as of November 2020; there were no phages between *Arthrobacter* Cluster FD phage Anjali (the 109th phage with 19,679 bp) and *Propionibacterium* Cluster BU phage Attacne (the 110th phage with 28,876 bp). Since this was a large gap of 10,000 bp, we set 19,679 bp as the upper cutoff for “small” phages, resulting in 109 small phages that have genome sizes ranging from 14,270 bp to 19,679 bp (Figure 1B). Of the 109 phages, 48 phages were isolated on *Arthrobacter*, 49 on *Microbacterium*, 9 on *Gordonia*, 2 on *Streptomyces*, and 1 on *Rhodococcus* (Supplementary Table S1). Of note, despite more than half of the 3,222 sequenced phages on PhagesDB that have been isolated on *Mycobacterium*, none of the 109 small phages infect *Mycobacterium* (the smallest phage known to infect *Mycobacterium* is 38,341 bp), suggesting that host type may be correlated with phage genome size.

**Figure 1.**
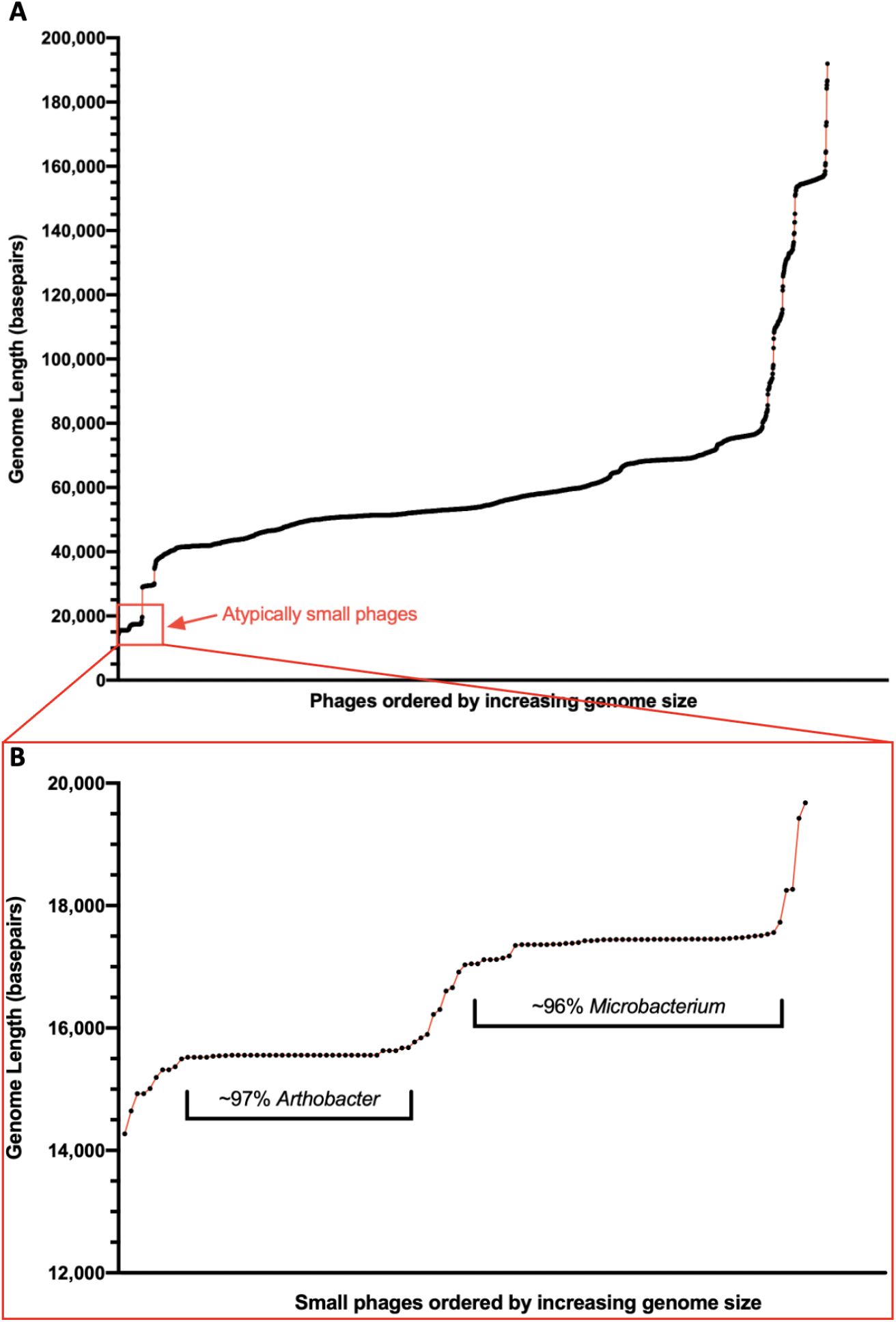
Atypically small phages primarily infect *Arthrobacter* and *Microbacterium*. **(A)** All 3222 sequenced phages (excluding draft phages) listed on PhagesDB in November 2020 are plotted by increasing genome size. **(B)** An enlarged graph of the atypically small phages region in panel A reveals 109 distinct phages, which primarily infect *Arthrobacter* and *Microbacterium*. Each black dot represents a distinct phage.

In order to better visualize and understand the genomic similarities and differences among the 109 phages, we selected 34 of those as representative phages. The 34 phages selected are highlighted in the list of 109 small phages (Supplementary Table S1) and compiled into a chart (Table 1) that summarizes general phenotypic characteristics of each phage, including cluster, host, morphotype, life cycle, and genome size. Our list of phages includes representatives from Clusters AN, AX, BO, *CW*, DM, EE, FD, FE, and GE that were isolated on *Arthrobacter*, *Streptomyces*, *Gordonia*, *Microbacterium*, or *Rhodococcus* hosts. The majority of the selected phages display *Siphoviridae* morphology and are lytic, except for Cluster BO *Streptomyces* phages that have *Tectiviridae* morphology [23] and Clusters CW and DM *Gordonia* phages that have a temperate life cycle.

### Nucleotide dot plot comparisons suggested minimal sequence conservation across the 34 representative phages

One method of determining the extent of relatedness within a group of phages is through the construction of a nucleotide dot plot, which allows qualitative comparison of nucleotide identity throughout the entire genome. In a comparison of the 34 representative small phages using this methodology, most observed similarity occurred between phages of the same cluster (Figure 2), which often have sizable regions of nucleotide identity [2]. Even then, certain phages from the same cluster such as the *Arthrobacter* phages from Cluster AX, *Gordonia* phages from Cluster CW, and *Arthrobacter* phages from Cluster FE which overall exhibited lower nucleotide conservation with other phages from their respective clusters.

**Figure 2.**
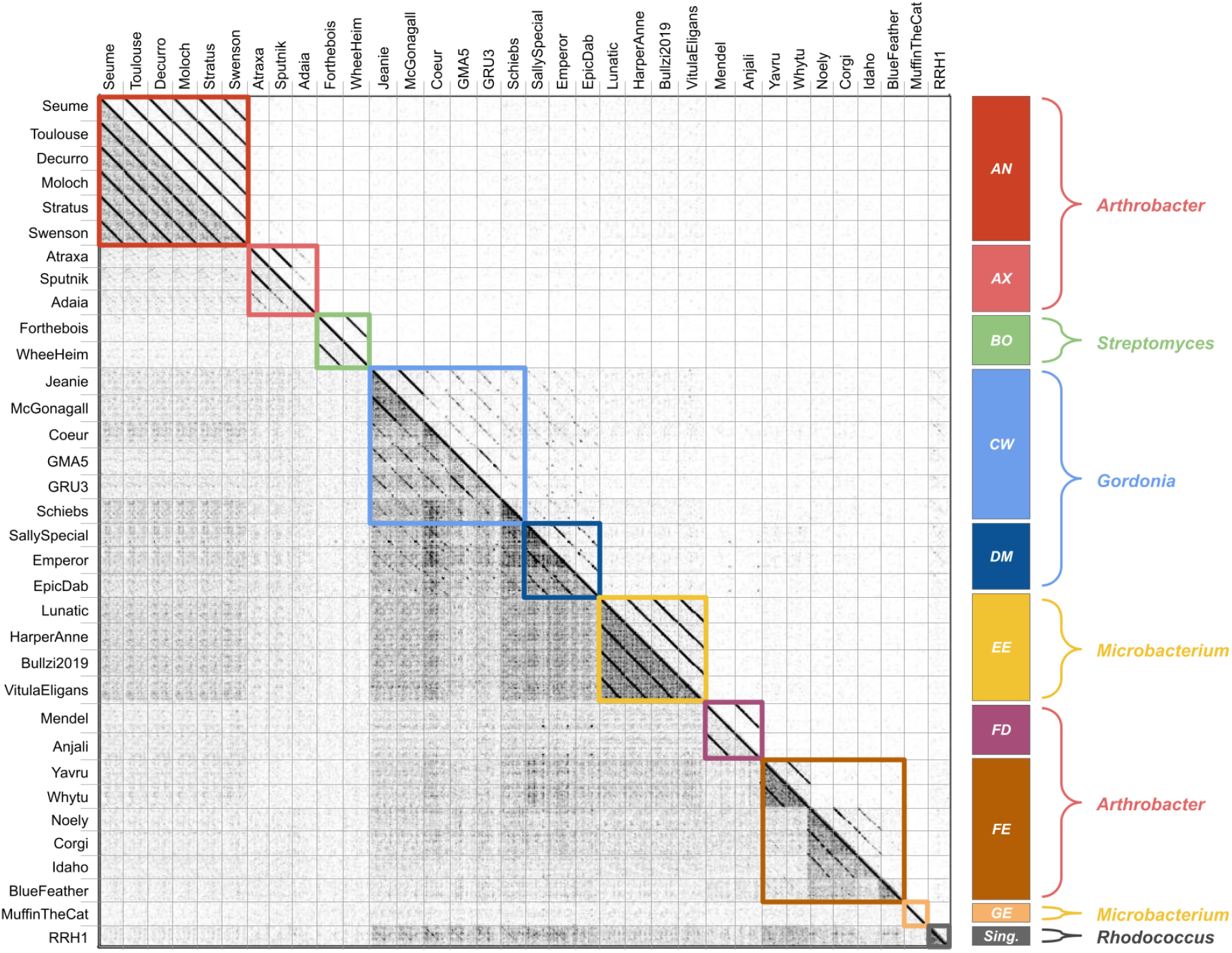
Nucleotide dot plot reveals minimal nucleotide sequence conservation among 34 representative small bacteriophages. Using the software Gepard 1.40, a dot plot was generated to observe alignments among the whole nucleotide genomes of 34 representative small bacteriophages with a word size of 12 on the right side, and word size of 10 on the left side. Minimal alignments were observed between phages of different clusters. The *Arthrobacter* Clusters AX and FE, as well as the *Gordonia* Cluster CW, contained select phages that had strong alignments to some but not all other members of their individual clusters.

### Amino acid dot plot and GCS comparisons exhibited greater levels of conservation, including significant similarity between *Arthrobacter* and *Gordonia* phages

Nucleotide comparisons can identify close relationships, but may miss more distant relationships between phages due to accumulated silent mutations and synonymous codon usage. A previous study found that *Arthrobacter* singleton BlueFeather, Cluster FE, and former Cluster FI phages shared low nucleotide similarity but considerable amino acid identity, resulting in the consolidation of these phages into Cluster FE. Contrary to the nucleotide dot plot analysis of small genome phages, dot plot analysis using whole genome concatenated amino acid sequences displayed multiple regions of amino acid identity not only between phages from the same cluster but also among phages from different clusters (Figure 3). For example, Cluster AN *Arthrobacter* phages exhibited similarity with Cluster AX *Arthrobacter* phages, and singleton *Rhodococcus* phage RRH1 was likewise similar to Clusters CW and DM *Gordonia* phages. These similarities were not apparent in the nucleotide similarity analysis.

**Figure 3.**
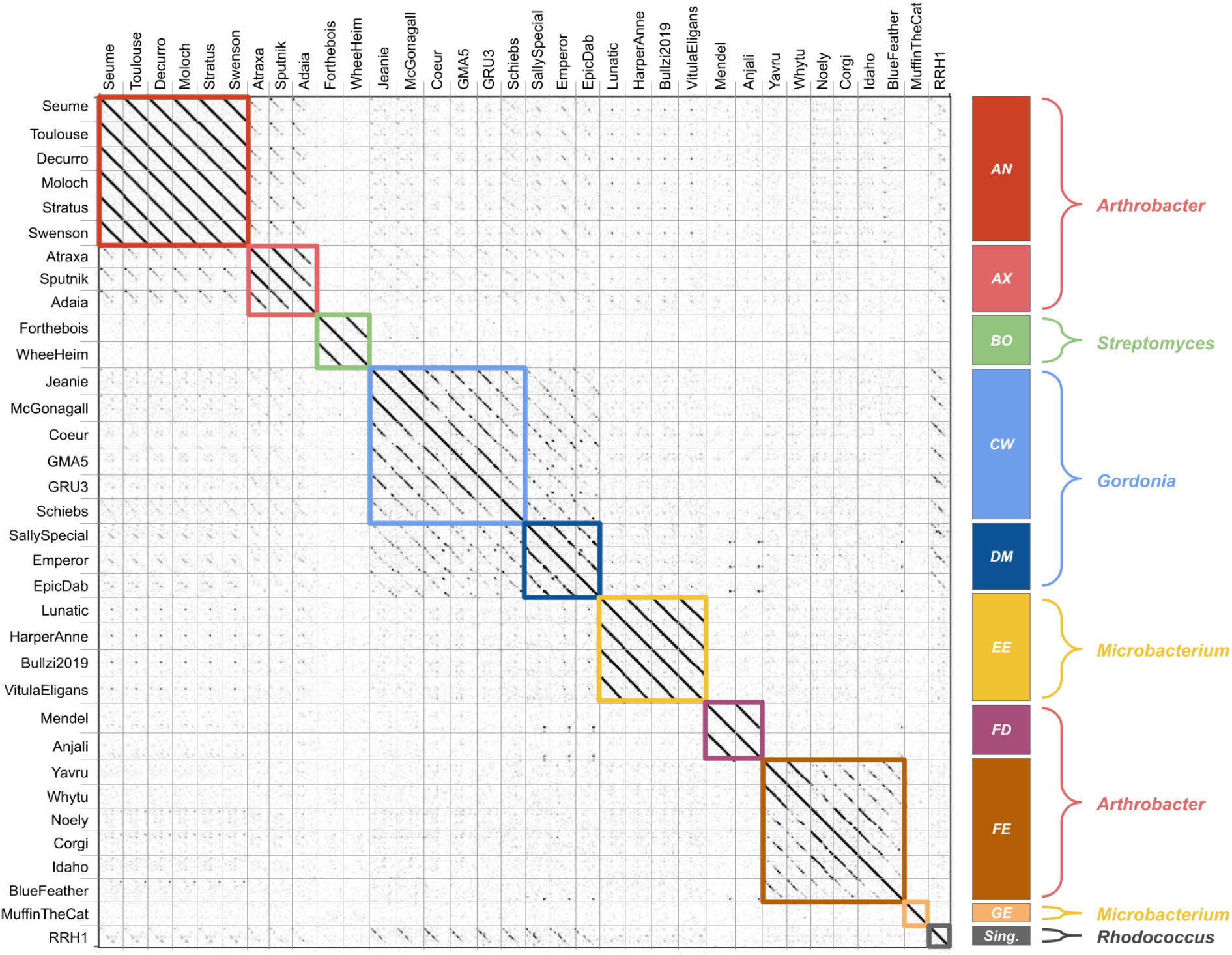
Amino acid dot plot exhibits substantial amino acid conservation among small phages. Using the software Gepard 1.40, a dot plot was generated to observe alignments among the concatenated complete amino acid sequences of 34 representative small phages using a word size of 5. *Arthrobacter* Clusters AX and FE, as well as the *Gordonia* Cluster CW, contained phages that all had moderate to strong alignments to all other representated phages within their clusters. Furthermore, alignments were observed between phages belonging to different clusters. Most notably, the *Rhodococcus* phage RRH1 exhibits significant alignment to many *Gordonia* phages in Cluster CW and DM.

To determine whether there were any phams that were shared between these small phages given the degree of amino acid similarity, we calculated gene content similarity among the 34 representative small phages. Consistent with the amino acid dot plot, significant GCS values (over 35%) were observed between phages from the same cluster as well as between Clusters CW and DM *Gordonia* temperate phages and Clusters AX and AN *Arthrobacter* lytic phages (Figure 4). Furthermore, singleton *Rhodococcus* phage RRH1 exhibited significant GCS values with Clusters AX and AN *Arthrobacter* and Clusters CW and DM *Gordonia* phages, further corroborating the amino acid identity seen in the dot plot (Figure 3). On the other hand, Cluster BO *Streptomyces* phages share the least amount of phams with other small phages — only displaying very low GCS values with Cluster GE *Microbacterium* phage MuffinTheCat. Although minimal nucleotide sequence similarity was seen between the small phages, our GCS analysis illustrates that the majority of small phages we sampled do share certain phams.

**Figure 4.**
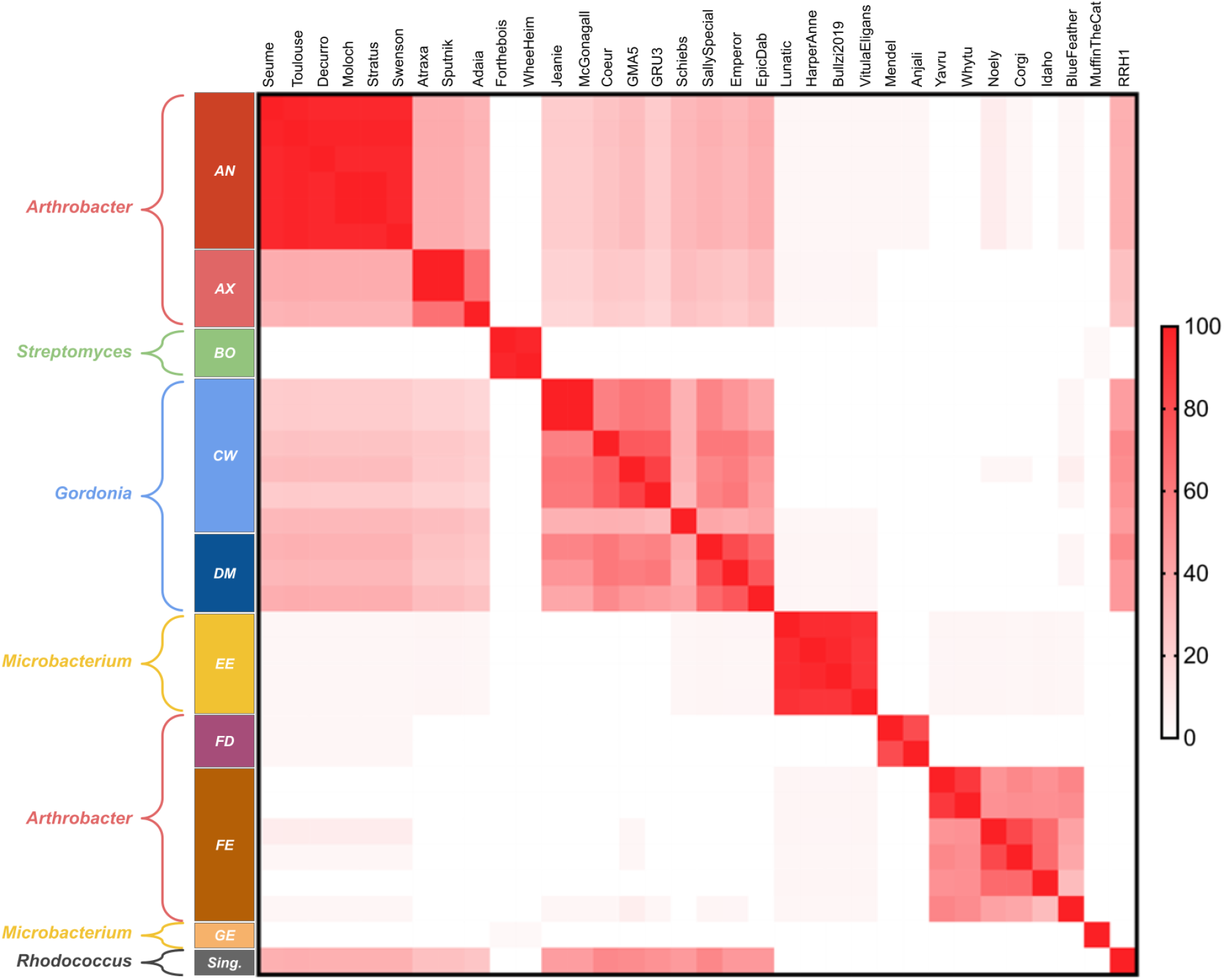
Shared gene content between 34 representative small bacteriophages. Using Prism9, a heatmap was generated to visualize pairwise GCS comparisons among all of the 34 representative small phages. Clusters CW and DM *Gordonia* temperate phages show significant GCS values (over 35%) with Clusters AX and AN *Arthrobacter* lytic phages. *Rhodococcus* phage RRH1 displays significant GCS values with Clusters CW and DM *Gordonia* phages and Clusters AX and AN *Arthrobacter* phages.

To better understand the subtleties of genetic relationships between these various small phage groups, a SplitsTree gene content network phylogeny was constructed for the 34 representative phages (Figure 5). *Arthrobacter* Cluster FE phages appear to have longer branch lengths from their most recent internal node than phages in the fellow *Arthrobacter* Clusters AN, AX, and FD, indicating that Cluster FE is one of the more diverse groups of *Arthrobacter* small phages, in correlation with its composition of phages that were originally in distinct clusters [24]. Likewise, the branches that include *Gordonia* Clusters CW and DM phages as well as singleton *Rhodococcus* RRH1 are highly diverse as well. In fact, based on the branches of this SplitsTree, the phages GRU3, GMA5, and Coeur (all subcluster CW2) and *Gordonia* phages McGonagall and Jeanie (all subcluster CW1) are more genetically similar to the *Rhodococcus* singleton phage RRH1 than to Schiebs (subcluster CW3). This is an instance in which phages from one cluster are more similar to non-cluster members than to phages within their cluster.

**Figure 5.**
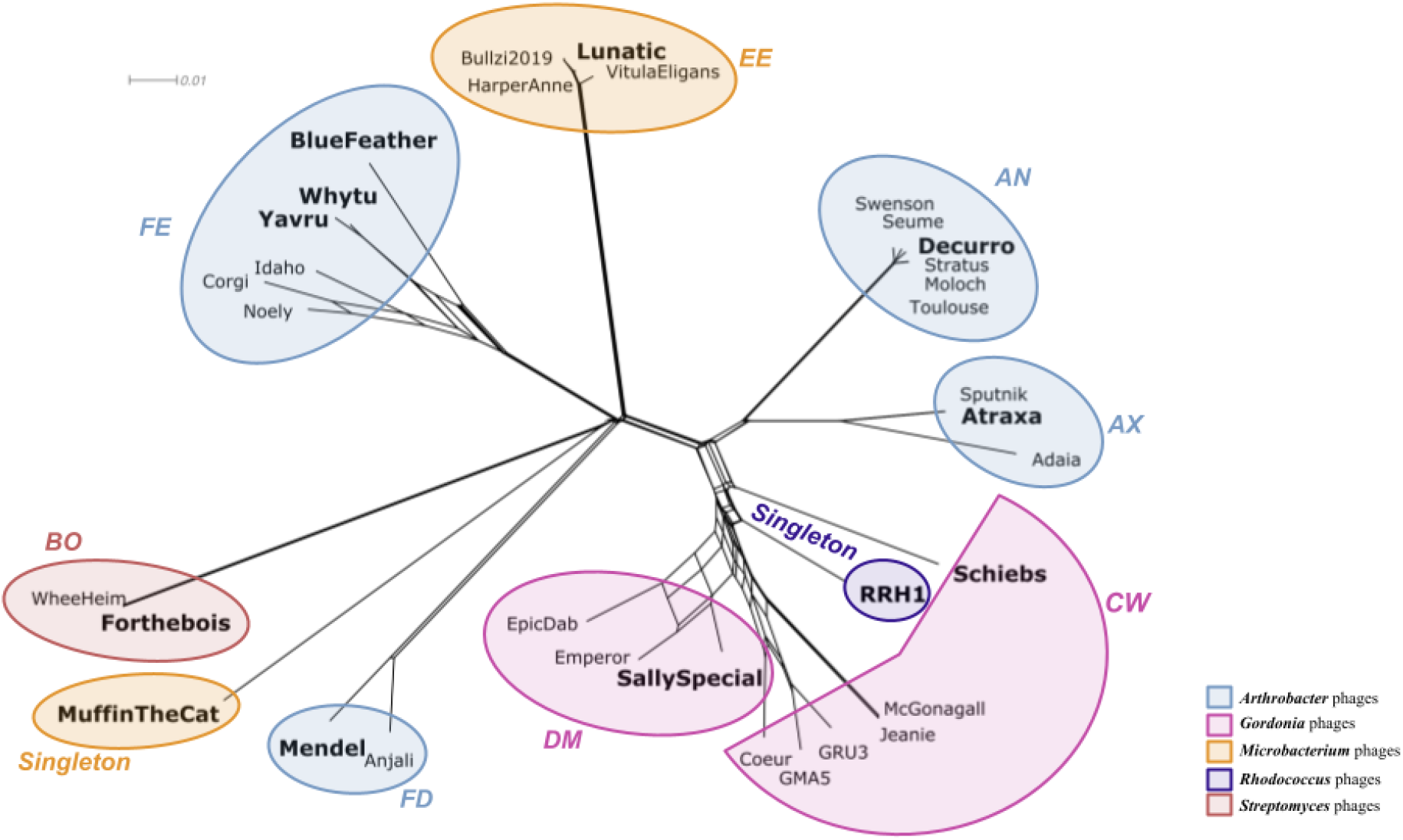
SplitsTree analysis unveils a continuum of diversity among small genome *Arthrobacter* and *Gordonia* phages. The phages in *Arthrobacter* Cluster FE have longer branch lengths than phages in the fellow *Arthrobacter* Clusters AN, AX, and FD, indicating that Cluster FE is one of the more diverse groups of *Arthrobacter* phages. The *Gordonia* phages GRU3, GMA5, and Coeur are part of the subcluster CW2 and phages McGonagall and Jeanie are part of the subcluster CW1; yet they are more genetically similar to the singleton RRH1 than to Schiebs, which belongs to the subcluster CW3. The bolded phages are representative phages for each cluster and appear in the pham map (Supplementary Figure S1).

### Pham maps suggested conservation of terminase and other genes among some small phages

As previous studies suggested the conservation of subunits of the same terminase phams in atypically small *Gordonia* phages [8], we performed a Phamerator map analysis to see if terminase and other additional genes would be conserved in other small phages as well. Indeed, the *Gordonia* small phages SallySpecial (Cluster DM) and Schiebs (Cluster CW) displayed conservation of the terminase phams. We noticed this two-unit terminase conservation not only among these atypically small *Gordonia* phages, but also with singleton *Rhodococcus* phage RRH1. Furthermore, these phages share significant GCS values, suggesting that despite their differing hosts, phage RRH1 is considerably similar to these *Gordonia* phages. Despite sharing no phams with any of the other phages in our Phamerator map analysis (Supplementary Figure S1), *Microbacterium* Cluster EE phage Lunatic shares its overall genome structure with the majority of other phages explored herein; all have a terminase protein, a portal protein, a capsid or protease fusion protein, and a tape measure protein, which can be expected given that these genes encode important functional components of the phage particle. Lunatic is not the only phage to exhibit similar genome architecture with small phages; other *Microbacterium* phages have also shared similar genome architecture to this group of atypically small phages [9].

## Discussion

The focus of this study was to characterize and better understand the intricate relationships between phages with small genomes. This was prompted by the isolation of phages Whytu and BlueFeather, both *Arthrobacter* phages that displayed extraordinarily small genome sizes. Our analyses of these 34 representative small phages, including Whytu and BlueFeather, showed minimal nucleotide similarity but greater amino acid and gene content similarity. Strong amino acid similarity in the absence of nucleotide similarity suggests that small actinobacteriophages have diversified enough that only distant relationships remain, perhaps due to gradual purifying selection of essential genes over their evolutionary history [25]. Phamerator map analysis of 11 of the 34 selected small phages further supports this point, as we observed intracluster conservation of phams between small phages that infect different hosts such as *Arthrobacter*, *Gordonia*, and *Rhodococcus*. The layout of genes and overall genome architecture were similar on an intercluster level, even across phages that did not share any phams, such as the *Microbacterium* phage Lunatic. This synteny in genomic architecture across otherwise distantly related phages has been observed in larger phages [9,26–28].

In our comparison of genome sizes and the isolation hosts of small phages, we observed that none of the 109 small phages were isolated on *Mycobacterium*, and that the smallest phage known to infect *Mycobacterium* was 38,341 bp long. This led us to ask several questions, such as why none of the 109 small phages in our study were isolated on *Mycobacterium*, and whether larger phages could have progressively evolved from smaller phages through direct acquisition of genes or through alternative mechanisms [29]. One study has suggested that some phams only present in one genome within a cluster may have been acquired by horizontal gene transfer [27], and another has suggested that larger phages may have progressively evolved from smaller phages through acquisition of genes via horizontal gene transfer [29]. Further research into an alternative hypothesis, such as whether reductive evolution in larger phages could have led to the loss of non-essential genes in small phages, would also provide more insight on the relationships between small and large phages.

Previous clustering methods have sorted phages with more than 50% nucleotide identity into the same cluster [27,28] and were later revised to utilize a 35% shared gene content parameter instead [8]. For instance, former Cluster FI phages and the singleton Phage BlueFeather have recently been combined with Cluster FE phages to form a single large cluster, Cluster FE. Therefore, it was expected that the branch lengths of the newly expanded Cluster FE in our SplitsTree analysis would be varied. On the other hand, branch lengths of the fellow *Arthrobacter* Cluster AN were extremely short and thus member phages were more similar. This variability in phage cluster diversity is observable in small phages across *Microbacterium*, *Gordonia*, and *Rhodococcus* hosts as well, and is an indicator of the continuum of actinobacteriophage diversity [8,9].

In a previous study, conservation of the large terminase subunits was observed among the atypically small *Arthrobacter*, *Gordonia*, *and Rhodococcus* phages studied [8]. Our Phamerator map analysis of 11 representative phages supported those findings, as the same terminase phams were present in *Gordonia* phages SallySpecial and Schiebs from Clusters DM and CW respectively. We furthermore noticed those same terminase phams present in the singleton *Rhodococcus* phage RRH1. Even more interestingly, we observed the same portal protein pham in the three *Gordonia* and *Rhodococcus* phages mentioned above in addition to the *Arthrobacter* phages Atraxa (Cluster AX) and Decurro (Cluster AN). These phages not only share these phams but also have a similar genome architecture with each other on a whole; this may be evidence for horizontal gene transfer that allowed these similar genes to be present in phages from a wide array of clusters and hosts [30]. This is perhaps unsurprising, as terminase and portal proteins tend to be more conserved than other genes [31], but these results are nonetheless valuable for further understanding the conservation of critical genes among small phages. Given the multiple different forms of evidence presented in this study through GCS, genome architecture, and conservation of phams, it seems that the *Rhodococcus* phage RRH1 may be more closely related to *Gordonia* phages than previously thought. While genomic similarity does not necessarily imply overlapping host ranges [32], we suggest host range experiments be conducted on RRH1 in light of its similarity to the *Gordonia* phages analyzed herein. It is possible that, since the *Rhodococcus* phage RRH1 shares so much in common with the *Gordonia* phages, it may be able to infect *Gordonia* hosts as well. The reverse situation may also be promising: to explore if the small *Gordonia* phages listed above are also able to infect *Rhodococcus*.

Overall, this study has defined a specific group of phages as so-called “small phages”: those with genome sizes that are under 20,000 base pairs and thus at least 10,000 bp smaller than every other actinobacteriophage sequenced to date on PhagesDB. We explored several patterns of conservation in their genomes among the various phage hosts and clusters represented. In addition, we verified low nucleotide identity but substantial amino acid and gene content similarity between some of these phages and also identified previously hidden relationships between the small *Rhodococcus* phage RRH1 and other small *Gordonia* and *Arthrobacter* phages, a testament to the complex dynamics that exist even among phages with such extraordinarily small genomes. This study only examined 109 small phages available on PhagesDB at the time, a mere handful of all the small phages that exist in the natural world. As more small phages continue to be collected and sequenced, further analysis of small phages will undoubtedly shed light on the concealed trends that define the nature of phage relationships.

## Acknowledgements

We thank Rebecca A. Garlena and Daniel A. Russell at the Pittsburgh Bacteriophage Institute for genome sequence and assembly and Travis Mavrich, Welkin Pope, Debbie Jacobs-Sera, and Graham Hatfull with the HHMI Science Education Alliance-Phage Hunters Advancing Genomics and Evolutionary Science (SEA-PHAGES) program for programmatic support. This research was supported in part by the Department of Microbiology, Immunology, and Molecular Genetics and the Dean of Life Sciences Division at UCLA. The authors acknowledge the use of instruments at the Electron Imaging Center for NanoMachines supported by NIH (1S10RR23057 to ZHZ) and CNSI at UCLA.

## Author Contributions

V.Y.T. and H.L. conceptualized the paper; V.Y.T., H.L., A.S.V., B.C.C., L.G.N., M.R., and S.N.F. performed experiments and drafted the paper; V.Y.T., H.L., A.K., A.C.F., and J.M.P. revised the paper; A.C.F. and J.M.P. supervised the research. All authors have reviewed and approved of this manuscript prior to submission.

This project was supported by the Microbiology, Immunology & Molecular Genetics Department and the Dean of Life Sciences Division at UCLA, with additional support for sequencing from the HHMI Science Education Alliance-Phage Hunters Advancing Genomics and Evolutionary Science (SEA-PHAGES) program.

## Supplementary Figures

**Supplementary Table S1.**
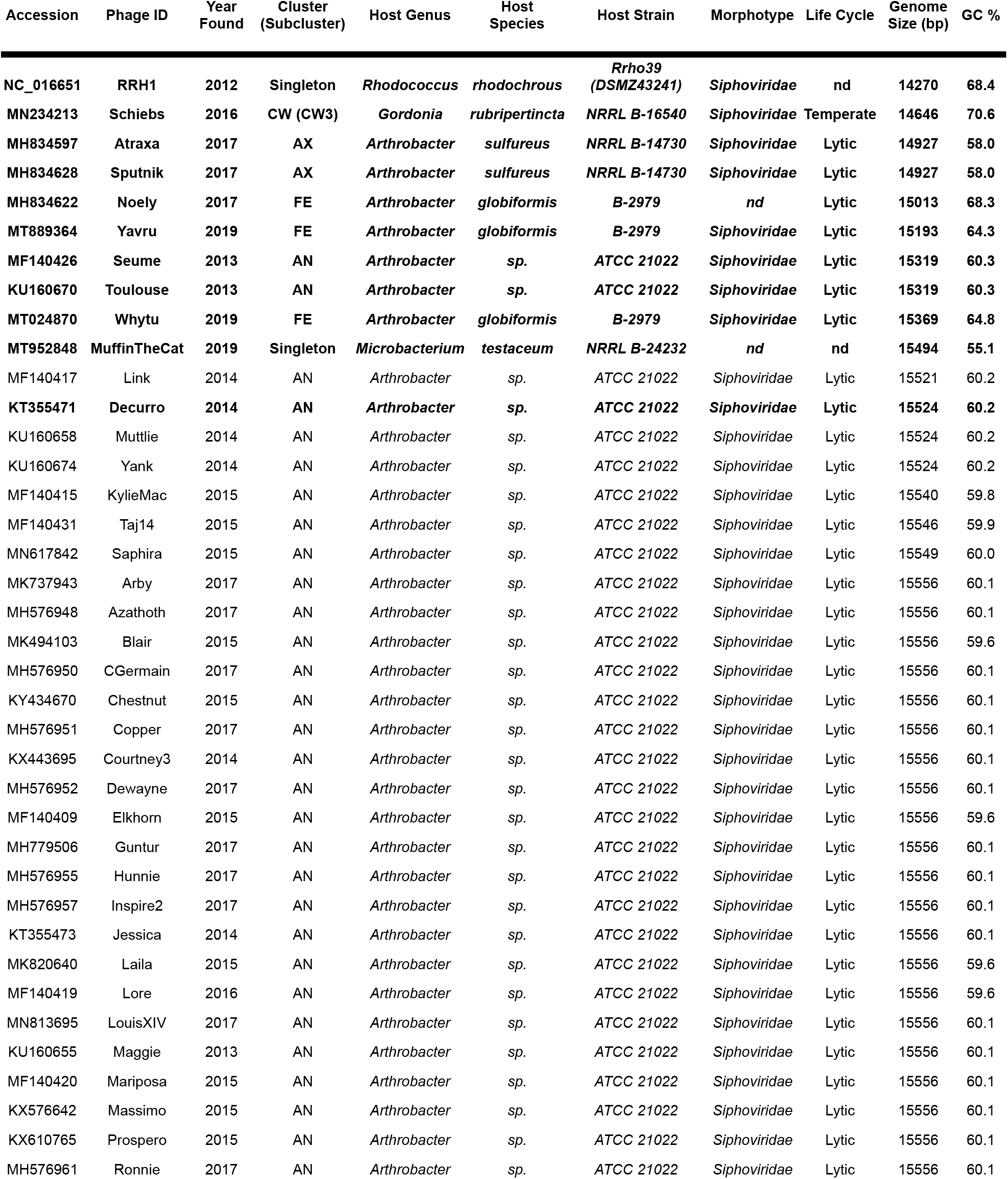
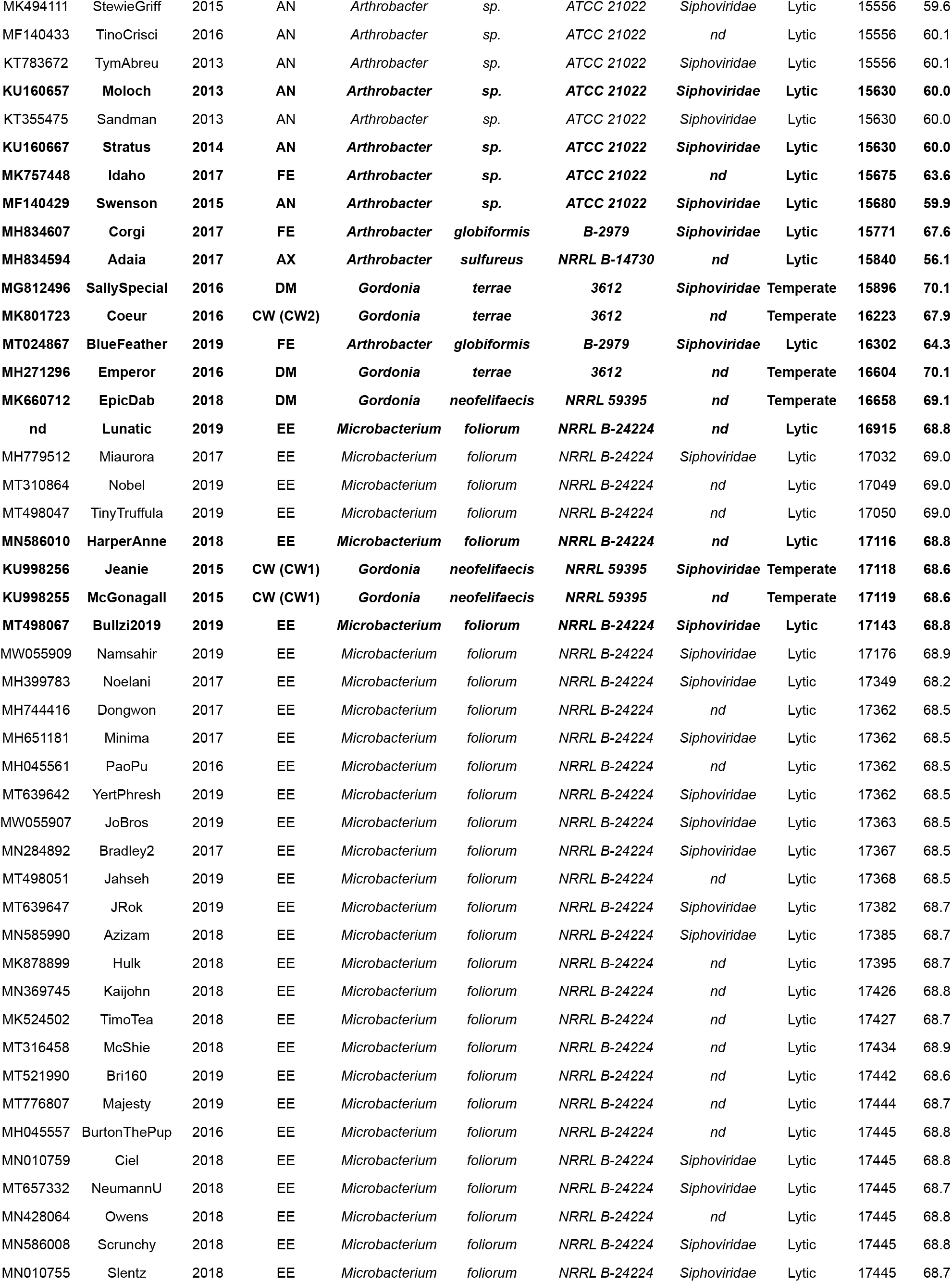
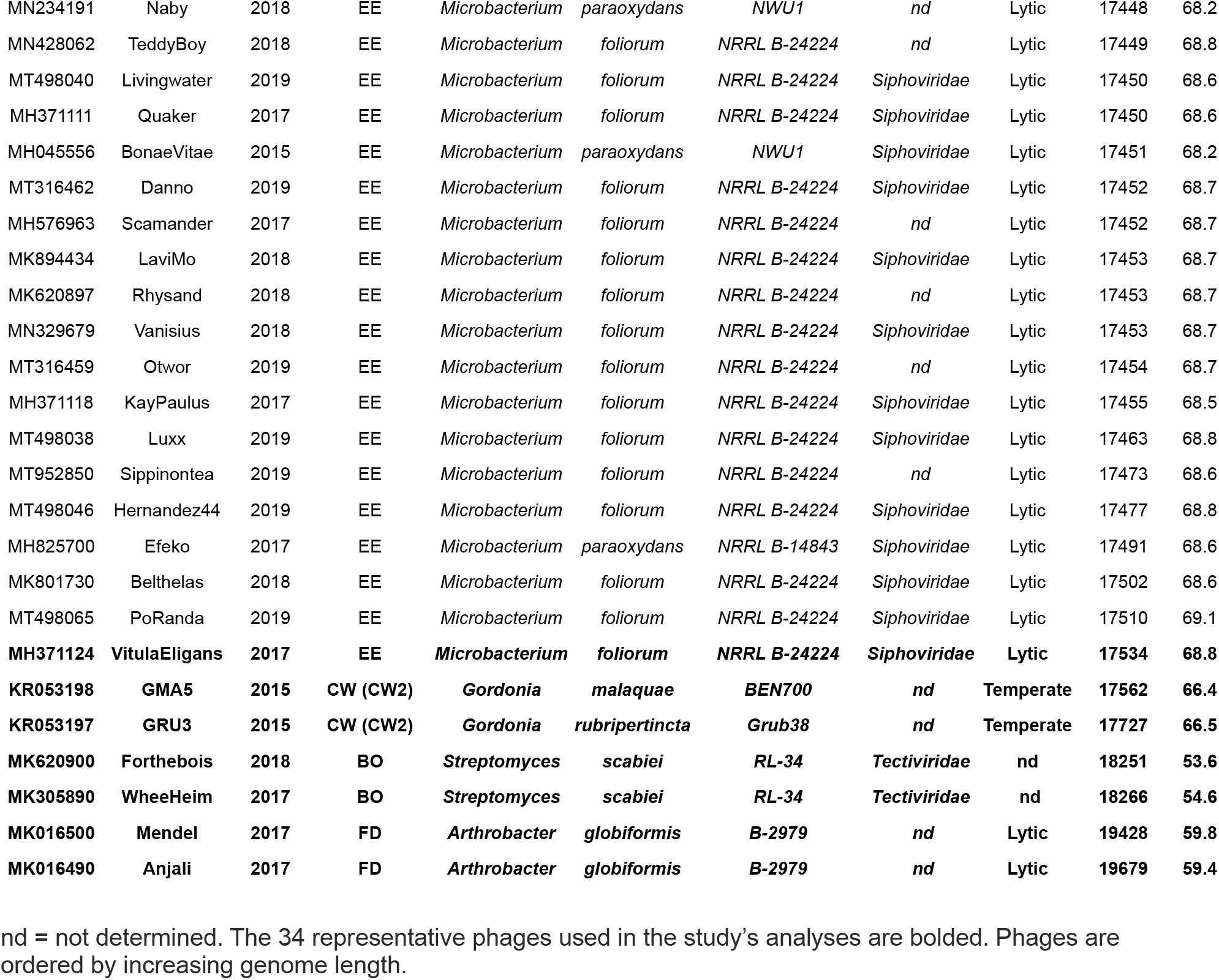
109 atypically small genome non-draft phages extracted from PhagesDB.

**Supplementary Figure S1.**
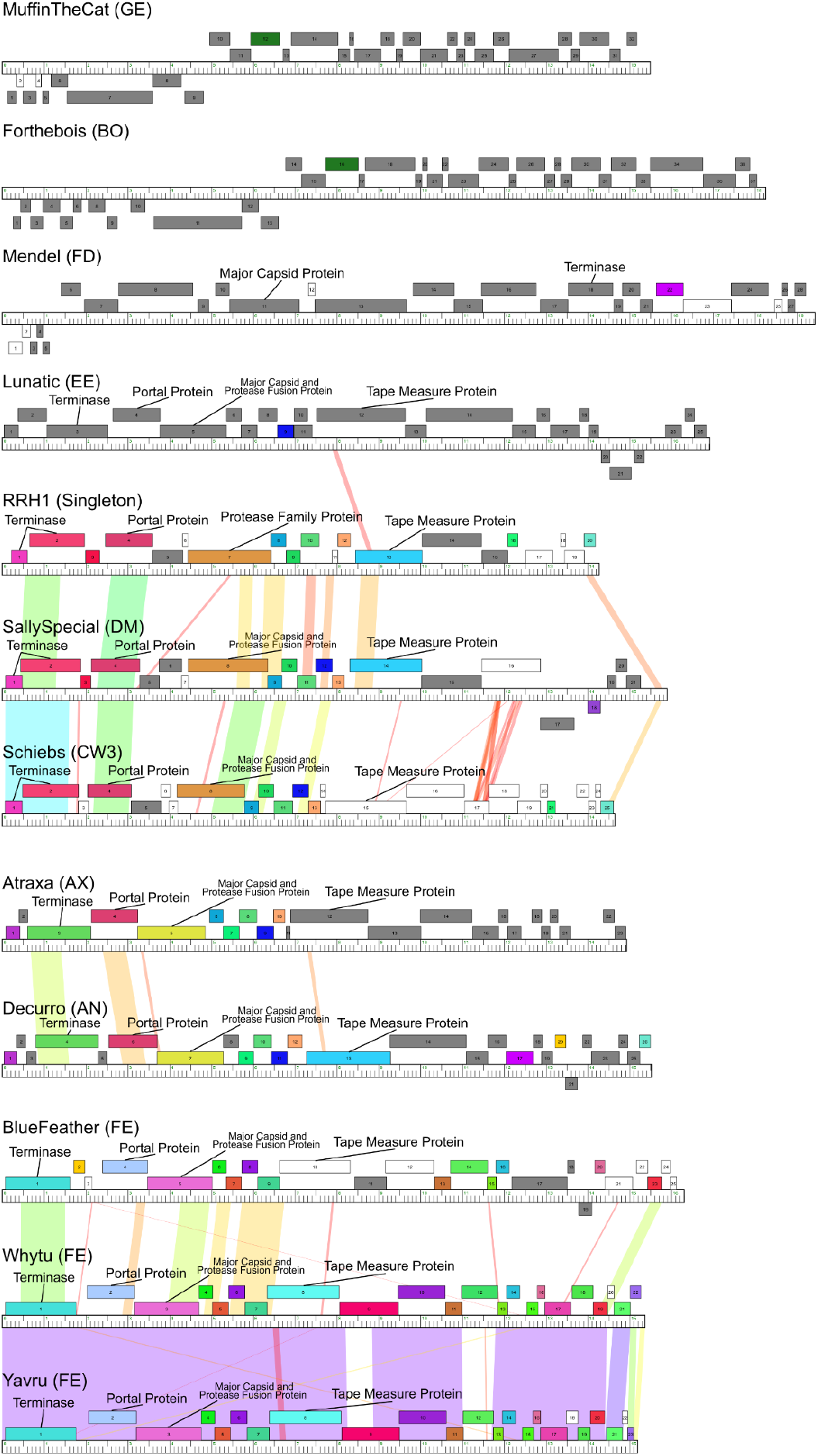
Pham maps reveal conservation of terminase and other phams among most small genome phages. 11 Phamerator genome maps, of at least one representative phage from each cluster within the group of 34 small genome phages, are displayed as of November 2020. Shared phams are colored; unshared phams are grayed out; orphams are uncolored. Shading indicates nucleotide similarity. Terminase conservation was observed in atypically small *Gordonia* phages: the group of phages that include RRH1 (Singleton), Decurro (AN), Schiebs (CW3), Atraxa (AX) and SallySpecial (DM) all exhibit intercluster conservation of the same terminase pham. All phages on this figure also share similar genome architectures; intercluster conservation of other genes besides terminase are also evident, such as the portal protein and tape measure protein.

